# Does temperature, light or inter-individual variation explain feeding motivation in a tropical freshwater predator?

**DOI:** 10.1101/2024.07.27.605438

**Authors:** Lucy J. Brown, Christos C. Ioannou

**Author notes:** Corresponding author: Christos C. Ioannou, School of Biological Sciences, Life Sciences Building, 24 Tyndall Avenue, University of Bristol, Bristol, BS8 1TQ, U.K. **Author Contributions:** L.B. and C.C.I. designed the research; L.B. performed the experiments; L.B. extracted the data from the videos; L.B. and C.C.I. analysed the data; L.B. and C.C.I. wrote the paper. **Competing Interest Statement:** The authors declare no competing interests.

## Abstract

Variability in environmental conditions in freshwater ecosystems are increasingly driven by human activity. Increased temperature and light intensity are among the anthropogenic stressors dramatically altering these ecosystems, for example through deforestation that reduces canopy cover of riparian vegetation. Simultaneous exposure to multiple stressors complicates predictions of responses to environmental stressors due to potential interactions, yet the interaction between temperature and light intensity on feeding motivation remains poorly understood. Here, a fully factorial design was employed to investigate the combined effect of increased temperature and light intensity on the feeding motivation of a freshwater predator, the pike cichlid *Saxatilia proteus*. Strikes toward food items were used to quantify the subjects’ motivation to feed. We found no effect of temperature or light intensity on feeding motivation, either individually or as an interaction. Our repeated measures design allowed us to test whether the predatory fish showed personality variation, i.e. consistent inter-individual differences, in their motivation to feed. While the time taken to make the first strike was not consistently different between individuals, the number of strikes in 1 minute, 3 minutes and the time taken for the 10^th^ strike (which were strongly correlated to one another), was consistently different between individuals. This variation could not be explained by variation in body length, which had no effect as a main effect. Variation between individuals is likely to be magnified in wild predators that are likely to vary more in experience and genetic background than the captive subjects in our study. Thus, we suggest that anthropogenic effects that alter the composition of individuals in a population of predators, for example selective harvesting, will have a greater ecological effect than direct short-term effects of variability in environmental factors.

## Introduction

Temperature and light are fundamental components of ecosystems and play a key role in driving species interactions by influencing foraging, and thus shaping ecological communities (Jägerbrand & Spoelstra, 2023; Kordas et al., 2011). However, as human activities continue to modify ecosystems and impose novel abiotic and biotic conditions, changes to inter-specific interactions may have complex impacts for community function and diversity (Guiden et al., 2019). Particularly in ectotherms, temperature increases energetic demand, which in turn impacts behaviour and physiology (Abram et al., 2017). Behavioural adjustments serve as the initial response to temperature fluctuations (Bailey et al., 2022), primarily manifested in changes to locomotor and feeding behaviours due to changes in the motivation to feed, allowing increased energy requirements to be met (Domenici et al., 2019; Volkoff & Peter, 2006). While the resilience of organisms to environmental stressors is heavily species- and context-dependent, tropical fish are likely to be disproportionately affected by warming because they possess narrow thermal windows and already live close to their thermal tolerance limit (Lapointe et al., 2018; Payne et al., 2016).

There is already evidence that increased energy consumption, due to elevated temperatures, is intensifying top-down competition and declining prey populations in the warming Indo-West Pacific Ocean (Johansen et al., 2015). This phenomenon can distort entire food webs and may already be affecting vulnerable freshwater ecosystems.

For animals that rely on visual cues, light is crucial for activities including reproduction, predator avoidance and foraging (Marchesan et al., 2005). In a predator-prey context, increased light intensity can aid predators locate prey (Fleming & Bateman, 2018; Richmond et al., 2004); conversely, it can make predators more conspicuous, inducing increased vigilance or fleeing behaviour in prey (Michels et al., 2024). Light in aquatic systems is predominantly influenced by surface illumination, depth and suspended particulates, which can change light intensity, colour composition and polarisation (Jägerbrand & Spoelstra, 2023). Since the development of human civilisation, deforestation has reduced the planet’s forests to less than 70% of their original extent (Bologna & Aquino, 2020). This is significant for freshwaters because the shading provided by riparian vegetation regulates both temperature and light conditions in freshwater habitats (Mosisch et al., 1999). In these ecosystems, which possess discrete physical boundaries, inhabitants may not be capable of large-scale movements in response to stressors from human-induced changes in environmental conditions (Morgan et al., 2001).

Many studies have researched the isolated effects of either temperature (Alfonso et al., 2021; Barbarossa et al., 2021) or light (Keep et al., 2021) in freshwater systems, controlling for other environmental stressors. To enhance the ecological relevance of research on responses to environmental stressors, there is an increasing focus on the effects of simultaneous exposure to multiple environmental stressors, which are ubiquitous in natural systems (Côté et al., 2016; Orr et al., 2020). Therefore, to form accurate predictions of their impacts it is essential to confirm how they interact as evidence suggests that they can combine in various complex ways (McFarland et al., 2012), which confounds projections of their net ecological impact (Thompson et al., 2018). Effects can be additive when the observed response is the sum of the responses from each stressor (Zanghi et al., 2024). Comparative responses are those dominated by the effect of only one stressor (Folt et al., 1999), whereas a ceiling effect may occur where the effect of an additional stressor is not evident (Ginnaw et al., 2020). An interaction between two stressors can be either synergistic (more than the sum of each response (Zanghi et al., 2023)) or antagonistic (less than the sum of each response (Ferrari et al., 2015)). Given this variability in outcomes, conducting empirical studies to assess the responses to stressors that co-occur is vital.

In addition to variation induced by changing environmental conditions, the past few decades have seen widespread interest in consistent variation in behaviour between individuals within populations, also known as animal personality variation, that cannot be explained by other traits such as age, sex and size (Dall et al., 2004; Sih et al., 2015). This research includes growing evidence that this consistent inter-individual variation can have ecological impacts (Brehm et al., 2019; Mittelbach et al., 2014); this is particularly the case regarding variation between individual predators in their motivation to feed, and hence the risk they pose to their prey. Consistent variation between individual predators has been documented particularly in predators in aquatic systems, demonstrating personality variation in direct measures of feeding, including the pike cichlid *Saxatilia frenata* studied *in situ* (Szopa-Comley, Duffield, et al., 2020), and northern pike *Esox lucius* (Nyqvist et al., 2012) and three-spined sticklebacks *Gasterosteus aculeatus* studied in the laboratory (Szopa-Comley, Donald, et al., 2020), although there were no consistent differences between individual blue acara *Andinoacara pulcher* tested under laboratory conditions (Szopa-Comley & Ioannou, 2022). Additionally, other studies have shown predators can vary in other behaviours that are likely to alter the risk they pose to prey, and hence the ecological impact of the predators, including activity (Nakayama et al., 2016), risk-taking tendency, i.e. boldness (Zhao & Feng, 2015), prey search behaviour (Patrick et al., 2014), space use (Villegas-Ríos et al., 2018) and foraging site fidelity (Harris et al., 2020).

The feeding behaviour of diurnal, predatory freshwater fishes is likely to be more severely impacted by warming and changes in the light environment than species capable of range shifts or those that are nocturnal which rely less heavily on visual cues (Freitas et al., 2021). Here, we test how the combination of increased water temperature and light intensity affects the feeding motivation of a freshwater fish, *Saxatilia proteus*, a species of pike cichlid (*Crenicichla*). Pike cichlids are piscivorous fish native to South American streams, rivers and lakes, relying mainly on visual cues to ambush prey (Szopa-Comley, Duffield, et al., 2020). In their natural habitat, pike cichlids will often experience elevated temperature and light intensity when changes in land use reduces canopy cover from removing riparian vegetation. Across river sites in the Northern Range mountains of Trinidad where *S. frenata* (which is closely related to *S. proteus* (Varella et al., 2023)) are abundant, Zanghi et al. (2024) demonstrated a positive correlation between temperature and light intensity, and declining light intensity with increasing canopy cover, although temperature and canopy cover were not significantly correlated (instead, temperature was negatively associated with flow rate). In our study, strikes on food items were used to measure feeding motivation and assess the subjects’ response to altered environmental conditions. Fish were exposed to control or elevated water temperature crossed with control or elevated light intensity in a fully factorial design to determine the individual and combined effects of these stressors. By using a repeated-measures design, i.e. with repeated tests per subject over a four-week period, we were also able to assess consistent inter-individual variation in the feeding response of these fish.

The study of Zanghi et al. (2024) also tested for the effects of naturally occurring environmental variation on the presence and predatory behaviour of piscivore fish in their study system. The presence of *S. frenata* was associated with warmer temperatures, and across the predators included in the study, predation pressure such as the number of attacks on the guppy prey presented as a stimulus also increased with temperature. We predicted that as the effects of temperature and light on visual predators are driven by different pathways, i.e. physiological for temperature and visual for light, that their effects would be additive rather than synergistic. Also based on field observations that presented live fish prey to wild pike cichlids (Szopa-Comley, Duffield, et al., 2020), we expected consistent inter-individual differences between the *S. proteus* in our study.

## Methods

### Experimental subjects and housing

Mixed-sex *S. proteus* (mean standard body length ± SD = 9.0 cm ± 1.0 at the time of testing) were reared at the University of Bristol, having been acquired from a commercial aquarium supplier. The 30 subadult (2-year-old) fish were housed individually in 45L tanks (L × W × H = 70 × 20 × 35 cm) enriched with sand and small stone substrate, plastic tube refuges and artificial foliage. Before the experiment, the water temperature was maintained at 25.8°C ± 0.3 SD, and the light regime followed a 12-hour light-dark cycle. For three weeks before testing started, subjects were exclusively fed 2ml of defrosted BCUK Aquatics Krill Pacifica once a day, which was used throughout the experimental trials because of its consistent sinking behaviour when introduced to the tanks. Opaque white plastic dividers were placed between and behind each tank to eliminate visual cues between subjects, and hence any influence between individuals during the trials.

During the trials, each tank had independent filtration provided by a Real Aquatics SF-101 internal sponge filter. The refuges, foliage and sponge filters were placed in the same configuration in each tank at the rear to ensure they would not obscure feeding during the filming of trials. Weekly water testing for nitrite, nitrate, and ammonia was conducted, in addition to 20% water changes on Mondays. Levels of nitrite and ammonia >0 ppm, or nitrate >20ppm, prompted water changes.

### Experimental treatments and protocol

Trials were conducted in the subjects’ housing tanks to minimise handling and stress, and hence facilitate and standardise feeding. Light intensity was manipulated using LEE 211 0.9ND 3-Stop Neutral Density Lighting Gel Filters to reduce lux levels within the tanks by approximately 70% (Table 1). The lighting filter sheets were cut to size (70 x 20 cm), with two slits at the rear to accommodate the air and water inflow tubes, enabling the lighting filters to sit flat on top of the tanks. The elevated light treatment comprised of the LED aquarium lighting without the filter (Table 1).

**Table 1:** Mean (± SD) temperature and light intensity in tanks during experimental trials measured by HOBO MX2202 loggers. Δ denotes the difference between control and elevated parameters. The values for the control and elevated temperatures is similar to the minimum and maximum temperatures recorded across sites in the Northern Range mountains of Trinidad that is habitat to *S. frenata* (Zanghi et al., 2024). However, light intensity from this study was typically much higher than in our study, with the minimum average light intensity recorded in the field being 111.6 lux.

|  | Temperature (°C) | Light intensity (lux) |
| --- | --- | --- |
| Control | 25.8 (± 0.3) | 45.5 (± 1.9) |
| Elevated | 29.1 (± 0.7) | 151.6 (± 38.3) |
| $\Delta$ | 3.3 (± 0.5) | 106.1 (± 36.4) |

AllPondSolutions and HDOM 25W aquarium heaters were positioned uniformly towards the rear of every tank; water temperature was manipulated by switching each of these heaters on (for the elevated temperature) or off (to maintain the initial housing temperature; Table 1). Each heater had a thermostat that was set to 29°C at the start of the experiment to ensure that minimal interference with the tanks would be necessary once the experiment started. 25W heaters were used in preference of greater capacity heaters so that the water temperature increased gradually to allow the fish to acclimate and not induce stress (Figure A1). The maximum temperature was limited to 30°C as this was similar to the highest recorded water temperature in the study of Zanghi et al. (2024). The average water temperature increase of 3.3°C in our study reflects projections for river warming over roughly the next 70 years (Liu et al., 2020).

Three HOBO Pendant MX2202 Waterproof Temperature/Light data loggers were used to record lux level and temperature at 5-minute intervals throughout the experiment, and were moved between tanks at the end of each testing day. In tanks assigned to the elevated temperature treatment and without a HOBO logger, the temperature was measured using an aquarium thermometer to confirm that the water was the correct temperature prior to testing. Experimental trials were conducted between 11 am and 4 pm, Tuesday to Friday over February and March 2024 for four weeks. As the fish were not fed on Saturdays and Sundays as part of their routine husbandry, they were fed on Mondays and experimental trials were only conducted Tuesday to Friday to standardise hunger across the testing days.

The four treatments consisted of a control, an elevated light treatment, an elevated temperature treatment and an interaction treatment where both light and temperature were elevated. The fully factorial design allowed for testing the effects of temperature and light intensity both independently and in combination. During testing, each individual experienced each of the four treatments once for one week by manipulating the temperature and light intensity in their home tank. Each tank was part of a block of 6 adjacent tanks; the order of the treatments for each tank was randomised, on the condition that each of the four treatments appeared at least once but not more than twice in that block of 6 tanks in a given week. On each day of testing, the order in which the blocks of 6 tanks were tested was randomised, as was the order of testing of the tanks within each block.

On the Friday of the week prior to the start of experimental trials, treatments for the first week were set up, involving the switching on of heaters and installation of light filters depending on which treatment each tank was assigned to, thus allowing the subjects time to acclimate to their respective treatments for three days before testing. Water testing was conducted each Monday morning in addition to water changes, ensuring a full 24-hour period elapsed before testing commenced the following day to allow the water temperature to reach the required temperature after the water change (Figure A1). After testing was completed on Fridays, the next treatment was set up for each tank for the following week.

Before the start of trials each day, 60g of BCUK Aquatics Krill Pacifica was defrosted in 20ml of filtered water. The trials were filmed using a Logitech C920 HD Pro Webcam mounted to a Manfrotto camera bracket and clamp. The camera was positioned centrally, facing the narrower vertical wall of the tank (i.e. the wall of dimensions 20 cm wide × 35 cm high), approximately 40cm in front of the bottom of the tank, angled at 40° upwards to be able to view the entire water column. Video recording was via QuickTime Player (version 10.5) at 1280×720 resolution and 30 frames per second. Recording was begun and 2ml of krill was injected using a 5 ml plastic syringe into the tank through a circular hole in the plastic lid of each tank (the hole was 2 cm in diameter and its centre 4 cm from the front of the tank, positioned centrally along the tank’s width). Feeding was recorded for 5 minutes from the addition of the food. The camera was then moved to the next tank in the testing order and the trial procedure repeated. A total of 480 trials were conducted; of these, data from 10 trials was missing due to malfunctioning of the recording software. In instances where the heaters malfunctioned, these trials were included as additional replicates for the non-elevated temperature treatments; this occurred in 9 trials across the duration of the experiment. After testing on the final day, each fish was caught in a net and their standard body length was measured using callipers.

### Data processing

Video recordings were analysed using the event-logging software BORIS (Friard & Gamba, 2016). *Crenicichla* make exaggerated jaw movements during feeding (Martinez et al., 2018), and this strike action was recorded as a point behaviour in BORIS. Strikes were used as a measure of feeding motivation rather than food consumption as the food items were not always visible to the experimenter in the video footage. The time in the video that each strike occurred within the 3 minutes after the food was added was recorded. A single experimenter logged the strikes in BORIS to avoid inter-experimenter variability. Four response variables were then calculated: the latency to the first strike (seconds), the latency for the 10^th^ strike (seconds), the number of strikes in 60 seconds, and the number of strikes in 180 seconds. The latency to the first strike was defined as the time from the introduction of the food into the tank until the first strike was made. Similarly, the latency for the 10^th^ strike was measured from when the food was added to when the 10^th^ strike occurred. For the number of strikes within 60 and 180 seconds, the time interval (60 or 180 seconds) started from the time of the first strike.

### Statistical analysis

R version 4.3.3 was used to conduct the statistical analyses. Using the lmer function in the lme4 package (Bates et al., 2015), each response variable (i.e. the latency to first strike, latency for the 10^th^ strike, number of strikes in 60s and number of strikes in 180s) was analysed separately in a linear mixed model (LMM). The temperature treatment (control or elevated) and light treatment (control or elevated) were included as fixed factors, as well as the interaction term between them. Testing day within the week (1 to 4), week number (1 to 4), and standard body length were included as main-effect only covariates in the models; these were all transformed using the scale function in R to avoid problems with model fitting. Consistent individual variation between the subjects was modelled using a random effect of subject identity. The DHARMa package (Hartig, 2019) was used to check the assumptions of normality of residuals (confirmed using Q-Q plots) and homogeneity of variances (confirmed by plots of the residuals vs. fitted values). As the latency to the first strike and the latency for the 10^th^ strike did not meet these assumptions, they were log-transformed and 1/square root transformed before analysis, respectively; these transformations were determined using Box-Cox transformations (Box & Cox, 1964), and the models met the assumptions after these transformations.

To determine whether light intensity, temperature and their interaction were important in predicting feeding motivation, we constructed models that differed in which of these explanatory variables were included, and compared these to one another and a model that lacked these variables (Tables 2, 3, A1 and A2). Model comparisons were carried out with the Akaike information criterion, corrected for small sample sizes (AICc), using the ICtab function from the bbmle package (Bolker & R Development Core Team, 2017). The model with the lowest AICc is considered the most likely model, and a difference of >2 AICc units between models can be considered strong support for the model with the lower AICc. If models are within 2 units of the most likely model, the model with the fewest parameters is favoured as being the most likely (Burnham & Anderson, 2002). The explanatory variables important in explaining variation in the response variable are those included in the most likely model. In these model comparisons, we also included models that tested whether body length and subject ID were important to include in the models. This was done by constructing models that removed body length from the model that lacked light and temperature, removing subject ID as a random effect from the model that lacked light and temperature, then removing both body length and subject ID from this model. Day within week and week were included as scaled covariates in all models. To quantify interindividual differences in feeding motivation, estimates of repeatabilities and their 95% confidence intervals (CIs) were obtained using the rpt function from the rptR package (Stoffel et al., 2017) for each of the four response variables.

**Table 2:** ΔAICc (difference in the Akaike information criterion, corrected for small sample sizes, between the model and the most likely model) model comparisons to determine which explanatory variables and the random effect of subject ID affected the latency to the first strike (natural logarithm transformed). The df is the number of parameters that are estimated in each model. All models include day of the week (1-4) and week (1-4) as main effects, and the random effect of individual identity unless otherwise stated. SBL is standard body length. N = 469 trials.

| Response variable: log(Latency to 1st strike (s)) | $\Delta AICc$ | df |
| --- | --- | --- |
| Subject ID random effect only | 0 | 5 |
| No random effect | 0.1 | 4 |
| SBL | 1.9 | 6 |
| SBL, no random effect | 1.9 | 5 |
| Temperature + SBL | 3.4 | 7 |
| Light + SBL | 3.9 | 7 |
| Temperature + Light + SBL | 5.4 | 8 |
| Temperature × Light + SBL | 7.4 | 9 |

**Table 3:** ΔAICc model comparisons to determine which explanatory variables and the random effect of subject ID affected the number of strikes in the 60 seconds after the first strike. See Table 2’s legend for further details. N = 470 trials.

| Response variable: Number of strikes in 60 s | $\Delta\text{AICc}$ | df |
| --- | --- | --- |
| Subject ID random effect only | 0 | 5 |
| Light + SBL | 0.9 | 7 |
| SBL | 1.8 | 6 |
| Temperature + Light + SBL | 3 | 8 |
| Temperature + SBL | 3.9 | 7 |
| Temperature × Light + SBL | 5.1 | 9 |
| No random effect | 149 | 4 |
| SBL, no random effect | 149.5 | 5 |

### Ethical note

The study was approved by the University of Bristol Animal Welfare and Ethical Review Body (UIN/23/074). Elevated temperature treatments were limited to 30°C and increased gradually between treatments (Figure A1) to minimise physiological stress. Water testing and changes were conducted weekly to ensure high water quality. After the experiment, the fish remained housed in the University of Bristol’s research facility to be used in future experiments.

## Results

The four measures of feeding motivation varied in the extent that they were correlated with one another (Figure 1). The latency to the 10^th^ strike and the number of strikes in both 60 seconds and 180 seconds were more strongly correlated than any of these were correlated with the latency to the first strike, although there was a moderate correlation between the latency to the first and 10^th^ strikes. For all four measures, there was no evidence that differences in light levels and temperature affected the feeding motivation of the fish, as models with these variables (either as the single main effect, both as main effects or as an interaction term) were less likely (or were only marginally more likely) than simpler models that lacked these terms (Tables 2, 3, A1 and A2, Figures 2a, 3a, A2a and A3a).

**Figure 1:**
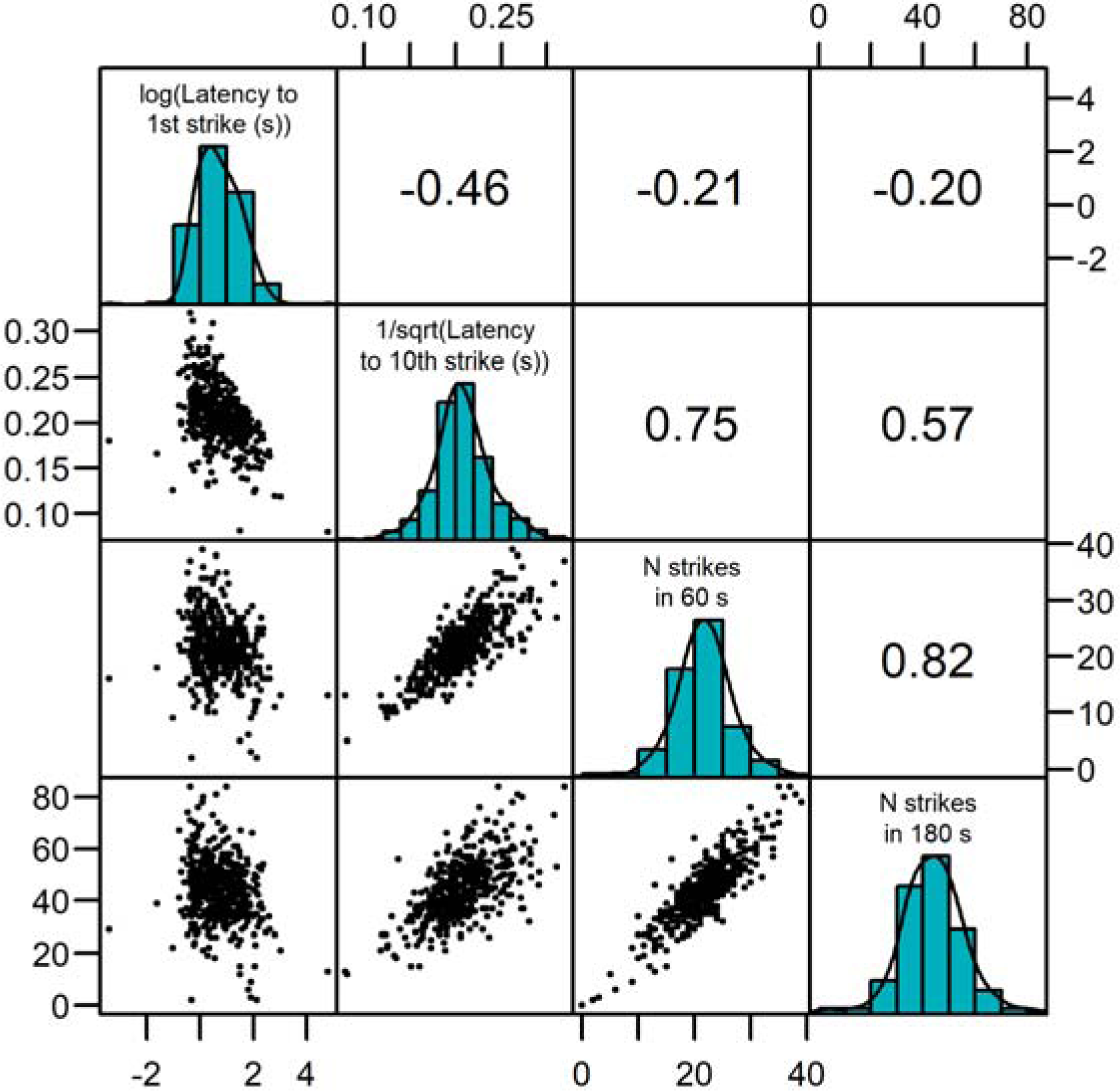
Correlations between the measures of feeding motivation (i.e., latency to the first strike, latency for the 10^th^ strike, number of strikes in 60s, and number of strikes in 180s). The values in the top right section of the plot are Spearman’s rank correlation coefficients. Note that with the 1/square root transformation for the latency to the 10^th^ strike, larger values indicate shorter latencies.

**Figure 2:**
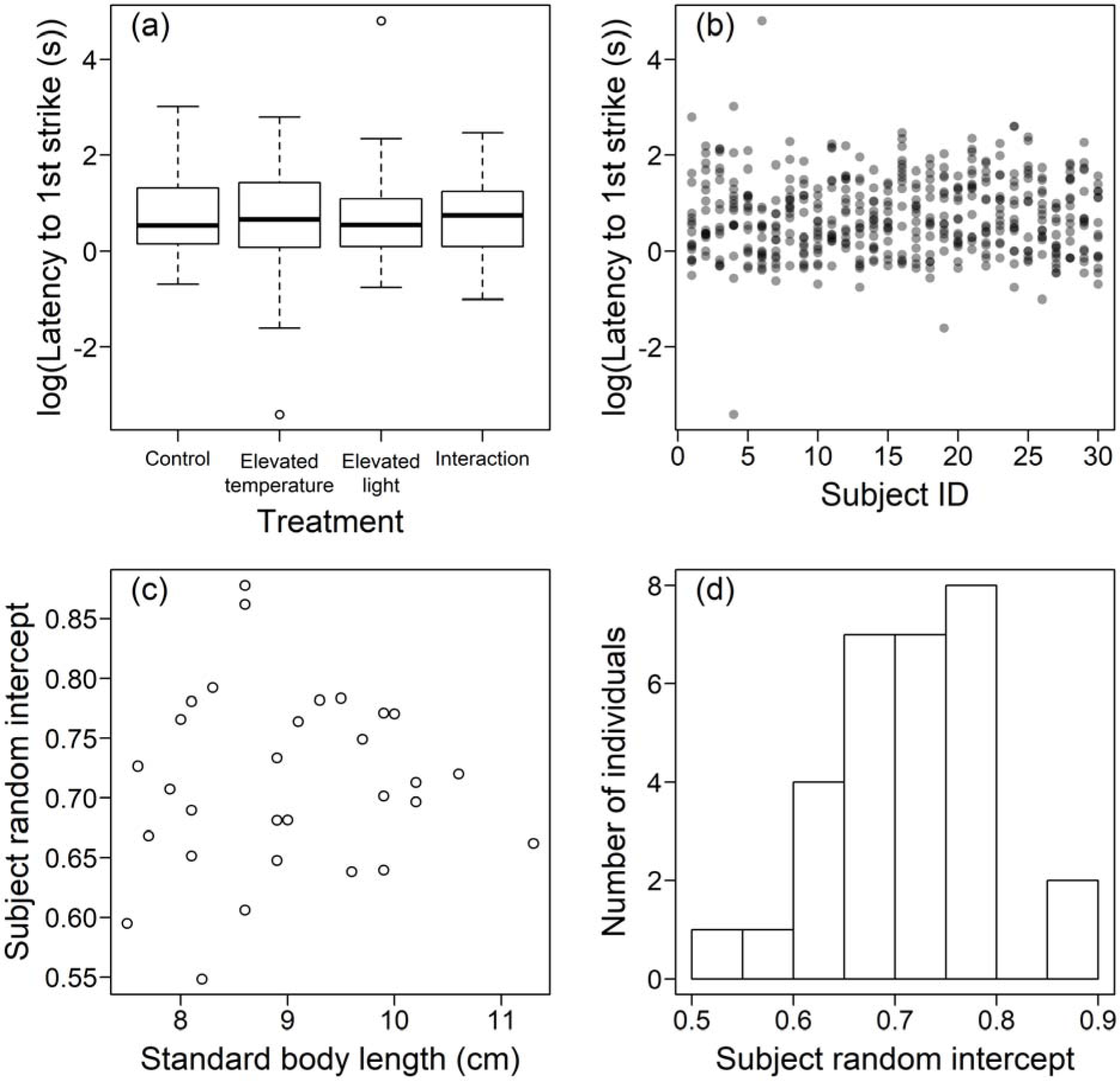
The latency to the first strike as a measure of feeding motivation. (a) shows how the latency to the first strike was affected by the manipulated environmental variables, and (b) how it varied between the different individual fish. Using the intercept fitted for the random effect of subject ID, (c) shows how differences between the fish in the latency to first strike varied with body size, and (d) shows the distribution of these random intercepts. In (a), the box length denotes the interquartile range, and the median value is represented by the horizontal black lines inside the boxes. The vertical black dashed lines indicate data points within 1.5 times the interquartile range above and below the upper (75%) and lower quartiles (25%). The circles represent outliers.

For the latency to the first strike, the model that lacked temperature, light and standard body length as explanatory variables, and the individual identity of the fish as a random effect, was 0.1 AICc units from the most likely model, suggesting none of these variables explained variance in the latency to the first strike (Table 2, Figure 2). The individual identity of the fish was, however, important to include in the models predicting the number of strikes within the first 60 seconds; the model with this as the random effect was the most likely model (Table 3). Figure 3b demonstrates the inter-individual variation in the number of strikes within the first 60 seconds. As the models with standard body length were less likely than the model with just the random effect, the consistent inter-individual variation was not due to differences in body size; Figure 3c demonstrates no association between body length and the individual-level intercepts fitted from the most likely model in Table 3. The distribution of the individual-level intercepts for the most likely model is unimodal (Figure 3d), suggesting that the consistent inter-individual variation was not driven by differences between sexes, as strong sex differences would be expected to generate a bimodal distribution (i.e. a mode for each sex). Consistent with the correlations between the measures of feeding motivation, these results were replicated with the number of strikes within 180 seconds and the latency to the 10^th^ strike (Tables A1 and A2, Figures A2 and A3).

**Figure 3:**
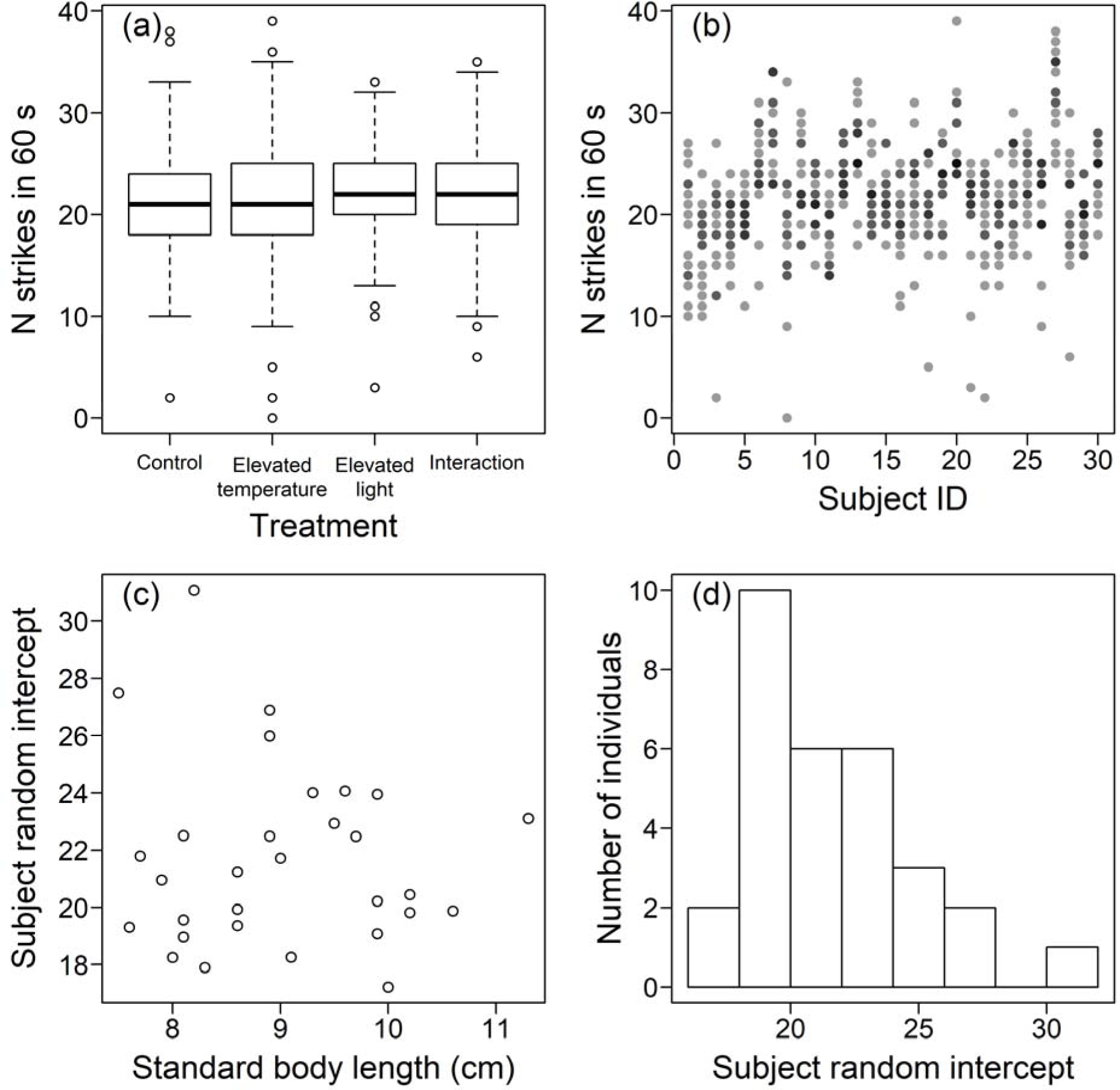
The number of strikes in the first minute as a measure of feeding motivation. Plotting as in figure 2. As the AICc model comparisons demonstrated strong consistent variation between the individual fish that could not be accounted for by differences in body size, and which are unlikely have been due to sex differences, we calculated repeatability estimates for the measures of feeding motivation. Consistent with the model comparisons, the repeatability estimate was low and the 95% confidence intervals included zero for the latency to the first strike (R = 0.03, 95% CIs: 0 - 0.082). However, the other three measures of feeding motivation demonstrated moderate repeatability estimates with confidence intervals that did not include zero (the number of strikes in 60 seconds: R = 0.38, 95% CIs: 0.234 - 0.505; the number of strikes in 180 seconds: R = 0.478, 95% CIs: 0.33 - 0.602; the latency to the 10^th^ strike: R = 0.26, 95% CIs: 0.147 - 0.382).

## Discussion

Interactions between multiple human-induced environmental stressors have been found to impose significant impacts across a range of behavioural contexts (Côté et al., 2016; Schmitz & Trussell, 2016). Although temperature and light intensity have been confirmed to impact feeding in other species (Domenici et al., 2019; Fleming & Bateman, 2018; Richmond et al., 2004; Volkoff & Peter, 2006), within the ranges tested in our study we found no effect of these stressors on the feeding motivation of *S. proteus*. There was, however, evidence of consistent inter-individual differences, i.e. personality variation, in this population of *S. proteus* in their motivation to feed. Thus, the composition of a predator population is more likely to impact prey, with potentially knock-on effects for the ecosystem via lethal and non-lethal effects (Binckley & Resetarits Jr., 2003; Resetarits Jr. & Pintar, 2016; Rudolf, 2012), than changes in temperature and light intensity that would be expected to occur through habitat change such as the removal of canopy cover over freshwater streams from deforestation.

The composition of a predator population with respect to their feeding motivation will vary depending on both natural and anthropogenic factors. For mesopredators such as pike cichlids, populations can become more risk-averse with an increased perception of predation risk, including a decreased feeding rate due to the trade-off between predation risk and foraging (Verdolin, 2006). In addition to this non-lethal effect, predation can selectively remove less risk-averse individuals from a population through direct consumption (Bell & Sih, 2007), although this is not always the case (Balaban-Feld et al., 2019). Harvesting by humans, either for recreation or commercially, is frequently non-random with respect to the risk-taking tendency of the individuals harvested, thus shifting the average risk-taking tendency in the population, usually toward shyer, more timid behavioural types (Arlinghaus et al., 2017; Biro & Post, 2008). Thus, harvesting of wild populations can act as an anthropogenic stressor indirectly impacting prey consumption by altering the behavioural composition of predator populations.

Increased temperature and light intensity are not the only effects of deforestation around freshwaters, which includes alteration of hydrological and water chemistry parameters (Castello et al., 2013; Ríos-Villamizar et al., 2017). Increased run-off and sedimentation caused by deforestation contributes to increased water turbidity, which declines the rate of predation by visual predators (Ehlman et al., 2020; Lunt & Smee, 2015). However, Zanghi *et al*. (2023) found that in turbid, warm water, guppies *Poecilia reticulata* (prey of the pike cichlid *S. frenata*) reduced shoaling and increased proximity to another predatory fish, the blue acara (*A. pulcher*), suggesting that these conditions could be advantageous to predators by reducing prey anti-predator behaviour.

Increased light penetration from reduced canopy cover from deforestation could be counteracted by more frequent and severe incidences of elevated turbidity, while the elevated temperature could increase the energetic demand of predators. Habitat change, driven by the single human activity of deforestation, can thus alter multiple environmental parameters that can potentially impact predator-prey interactions in freshwater ecosystems.

By conducting the trials before the fish were fed that day, our study was designed to infer feeding rates when motivation to feed would be high. However, over longer time scales, piscivore fish become satiated when prey are abundant, which is a likely driver for type II and type III functional responses being common in piscivores under natural conditions (Moustahfid et al., 2010). When prey are abundant, individuals with a greater motivation to feed will satiate earlier, reducing their consumption of prey, and reducing the difference in feeding rates compared to less motivated predators that are less sated. Such a state-behaviour feedback (Sih et al., 2015) will reduce inter-individual variation in feeding. Research on three-spined sticklebacks (*G. aculeatus*) has demonstrated consistency in individual differences was reduced in a foraging context but maintained in control trials where no food was present (MacGregor et al., 2021), although the opportunity to forage tended to make individuals less predictable rather than reduce variation between individuals. Further research in this area is needed to determine the extent of consistent inter-individual variation between predators at different prey densities. Another factor which may suppress consistent differences between individuals is conformity, where individuals become more similar in their behaviour due to signals or cues from one another (Ioannou & Laskowski, 2023).

Although adult pike cichlids are not found in social groups other than reproductive pairs, multiple individuals can occupy the same pools and be within visual range of one another (Szopa-Comley, Duffield, et al., 2020; Zanghi et al., 2024), and social information can be used in a foraging context even in fish species that do not live in groups (Webster & Laland, 2017). The extent to which consistent inter-individual variation within a predator population is supressed by social information leading to conformity deserves further study.

The latency to the first strike was not consistently different between individual *S. proteus*, and this variable was not strongly correlated with the other three variables used to quantify feeding motivation (the latency to the 10^th^ strike, and the food consumed in the first minute and first three minutes), which were all correlated with one another. While the latency to the first strike has been used as an indicator of feeding motivation in fish previously (Volpato et al., 2013), in our study this variable may have been sensitive to the location and orientation of subjects within the tank when the food was introduced, contributing unaccounted variation in the statistical models predicting the latency to the first strike. The feeding trials in our study were conducted in the fish’s home tanks to reduce stress by being a familiar environment and to avoid handling before the trials, and hence facilitate feeding. With a different tank arrangement that would allow filming from multiple perspectives, the fish’s location and orientation within the tank could be quantified and included as covariates in the statistical models (as in MacGregor et al., 2020). The use of pose tracking software (e.g. Pereira et al., 2022) would also allow kinematic measurements of the strikes.

Alternatively, when investigating the effect of temperature on feeding motivation in Pacific halibut *Hippoglossus stenolepis*, Stoner *et al*. (2006) arranged multiple vertical feeding tubes in each tank for food introduction. This setup allowed food to emerge at the greatest distance from the test subject, standardising subject positioning within tanks and minimising potential confounding effects on latency measurements. In our study, the variable location and orientation of the subjects within the test tank when the food was first introduced is likely to have had a reduced contribution to the variability in strikes after the first strike, explaining why we were able to detect consistent inter-individual variation in the latency to the 10th strike, and the food consumed in the first minute and first three minutes.

Overall, our study suggests that feeding motivation in pike cichlids is robust to relatively short-term exposure to changes in temperature and light intensity that is likely to be associated with removal of canopy cover from deforestation adjacent to freshwater streams. Instead, individuals of *S. proteus* show consistent differences in their feeding motivation, and these differences were unrelated to the body size of the individuals. A similar trend of consistent inter-individual variation in feeding that was independent of body size has been also demonstrated in another pike cichlid, *S. frenata*, under field conditions (Szopa-Comley, Duffield, et al., 2020), and in northern pike *E. lucius* (Nyqvist et al., 2012). Despite the prevalence of consistent inter-individual differences in predator populations, the ecological impacts of these differences have yet to be extensively explored, which are likely to be varied. For example, individual predators posing a greater risk to prey are likely to have different spatial distributions to less dangerous individuals, creating risk landscapes for their prey (Dammhahn et al., 2022; Steinhoff et al., 2020). Predator-prey interactions in freshwater aquatic systems, which can be studied under both laboratory and field conditions and are amenable to experimental manipulation as well as observation, are particularly promising for future studies in this area.

## Acknowledgements

This research was funded by a Natural Environment Research Council grant no. NE/P012639/1 awarded to C.C.I. We thank Dr. Martin J. How for advice on manipulating light intensity.

## Appendix

**Table A1:**
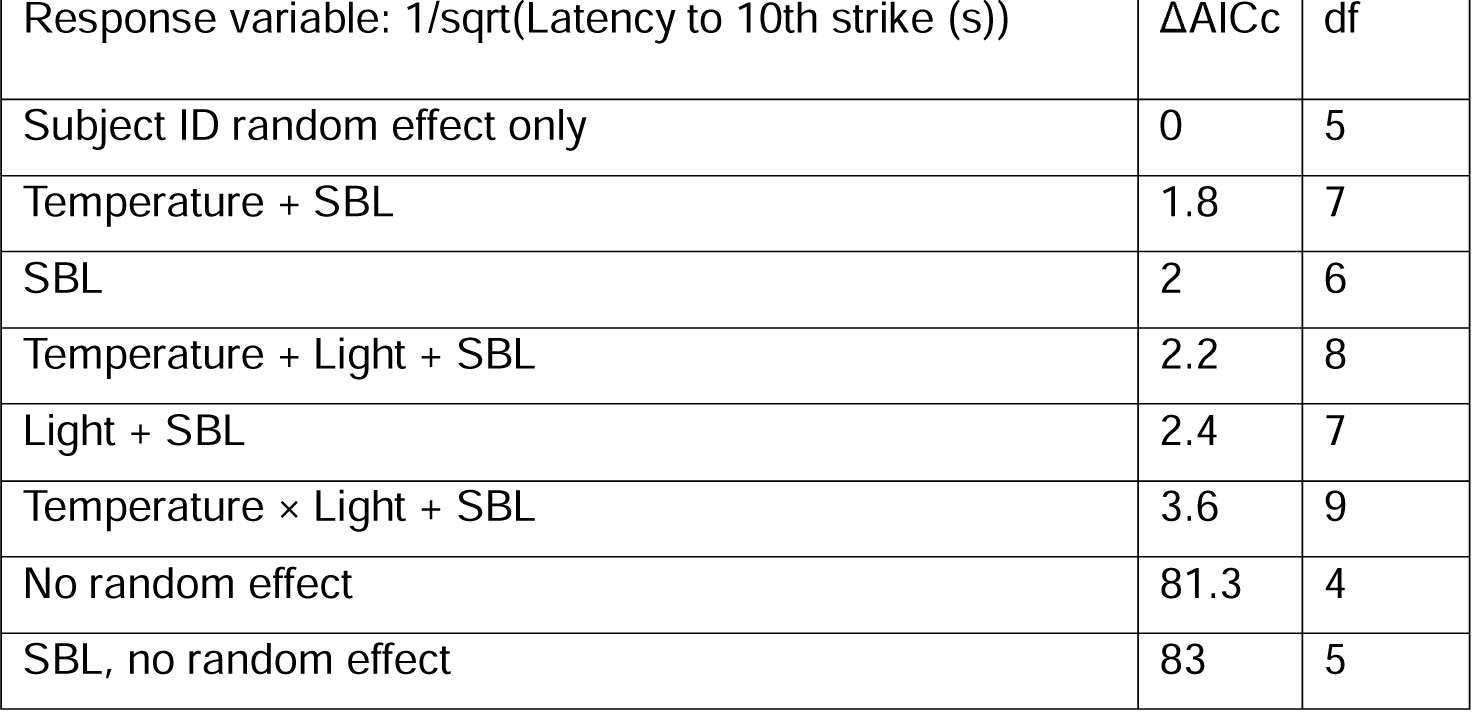
ΔAICc model comparisons to determine which explanatory variables and the random effect of subject ID affected the latency to the 10th strike. See Table 2’s legend for further details. N = 464 trials.

**Table A2:** ΔAICc model comparisons to determine which explanatory variables and the random effect of subject ID affected the number of strikes in the 180 seconds after the first strike. See Table 2’s legend for further details. N = 470 trials.

| Response variable: Number of strikes in 180 s | $\Delta AICc$ | df |
| --- | --- | --- |
| Subject ID random effect only | 0 | 5 |
| SBL | 1.4 | 6 |
| Light + SBL | 1.7 | 7 |
| Temperature + SBL | 3.4 | 7 |
| Temperature + Light + SBL | 3.6 | 8 |
| Temperature × Light + SBL | 4.6 | 9 |
| SBL, no random effect | 217 | 5 |
| No random effect | 220 | 4 |

**Figure A1:**
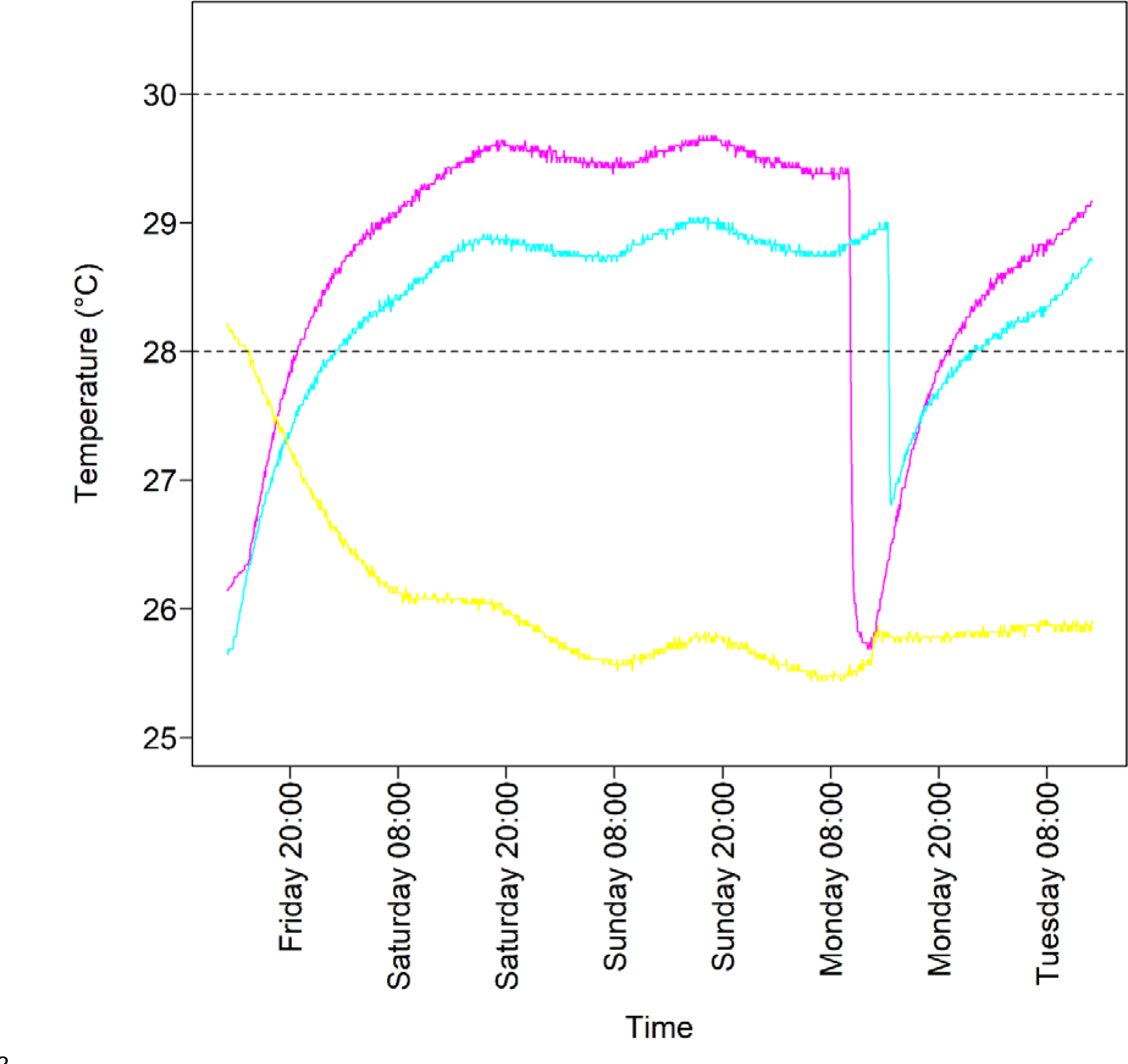
Water heating and cooling profiles for tanks 9 (magenta), 22 (cyan) and 28 (yellow) during the transition from week 1 to week 2 of trials. Temperature readings were captured by HOBO MX2202 loggers at 5-minute intervals. The black dashed lines at 28°C and 30°C represent the lower and upper thresholds for elevated temperature treatments. The transition from week 1 to week 2 treatments occurs on Friday, with trials commencing on Tuesday. The sharp temperature decline on Monday was due to the activation of the inflow system for water changes. Within 24 hours, the heaters in tanks 9 and 22 reached the elevated temperature range in preparation for the trials, while tank 28 remained at the control temperature.

**Figure A2:**
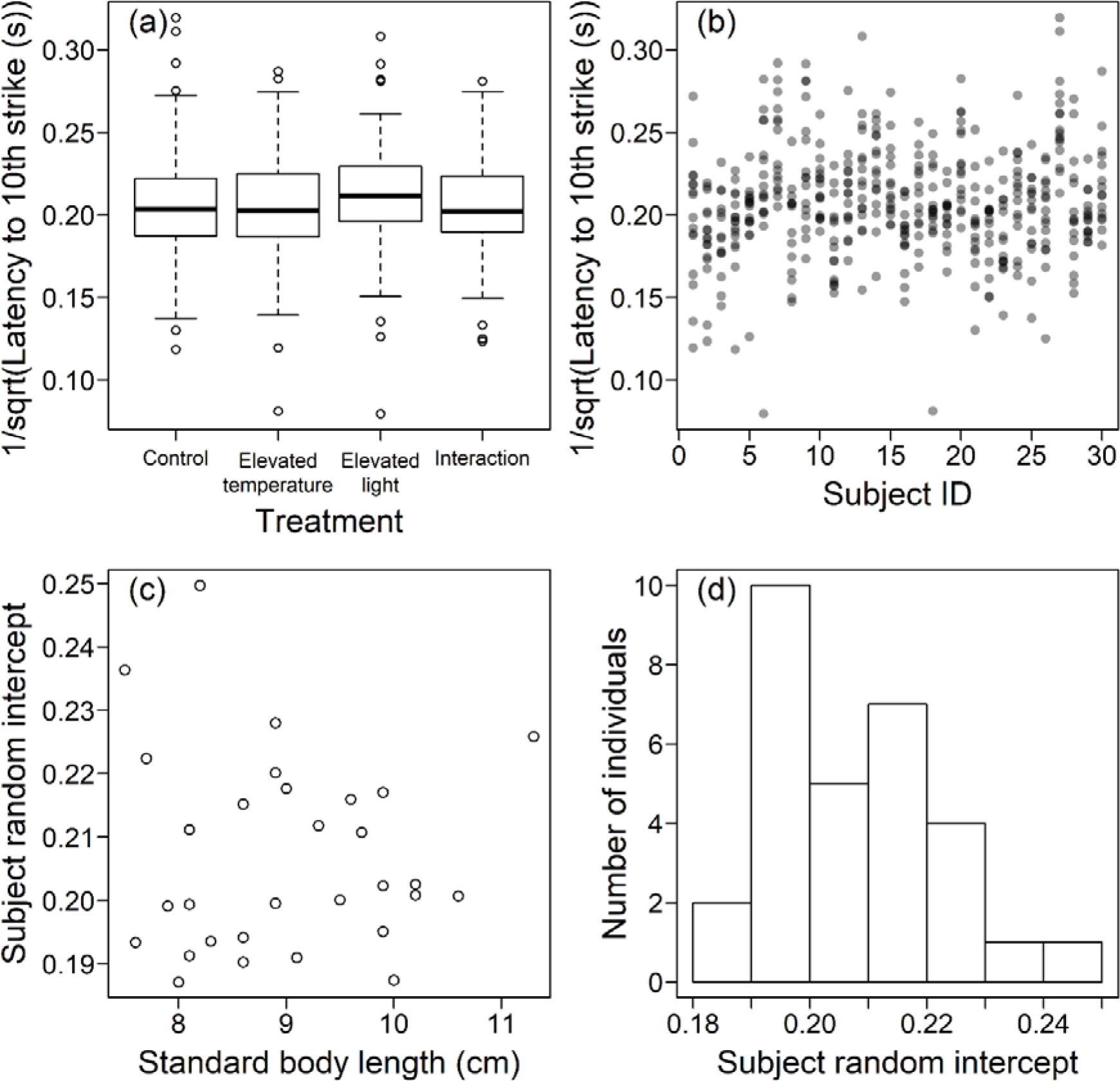
The latency to the 10th strike as a measure of feeding motivation. Plotting as in figure 2.

**Figure A3:**
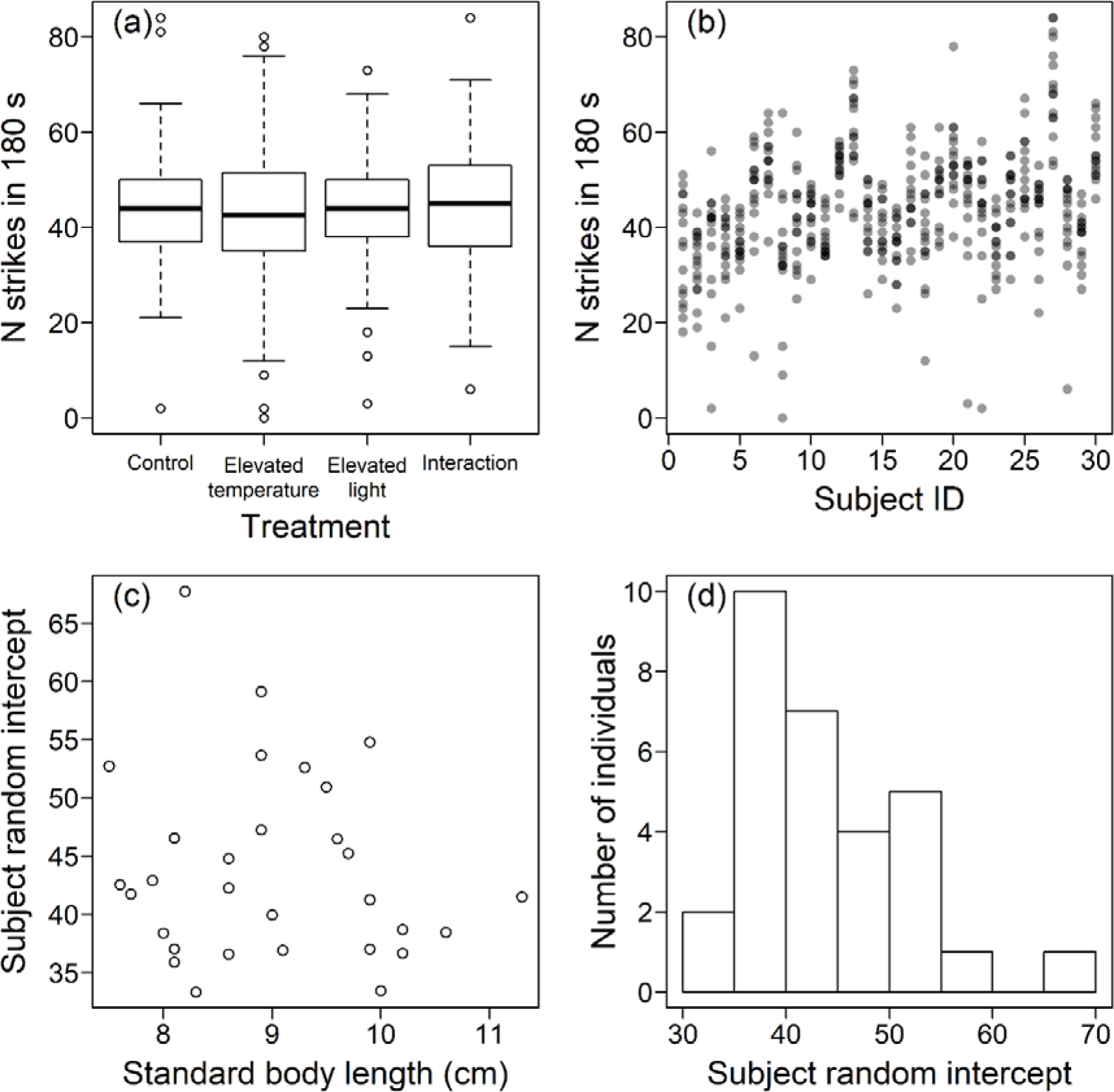
The number of strikes in the first three minutes as a measure of feeding motivation. Plotting as in figure 2.

## Notes

### Competing Interest Statement

The authors have declared no competing interest.

